# Functional Human Retinohypothalamic Tract Assembloid Model for Circadian Rhythm Research

**DOI:** 10.64898/2026.01.21.700761

**Authors:** Burak Kahveci, Ece Çakıroğlu, Hümeyra Nur Kaleli, Seden Nadire Harputluoğlu Efendi, Burak Derkuş, Ömer Faruk Özelçi, Sedat Nizamoğlu, İbrahim Halil Kavakli, Şerif Şentürk, Hayrunnisa Bolay, Bayram Yılmaz, Sinan Güven

## Abstract

The retinohypothalamic tract (RHT) is the primary pathway for circadian photoentrainment. Rodent models exhibit a significant translational gap for human physiology due to their nocturnal nature. To overcome this, we developed a functional human RHT assembloid by fusing human pluripotent stem cell (hPSC) derived retinal and hypothalamus organoids. Characterization revealed mature retinal ‘brush borders’ and the preservation of melanopsin-expressing intrinsically photosensitive retinal ganglion cells (ipRGCs) integrated via excitatory glutamatergic synapses. Multielectrode array (MEA) analysis confirmed synchronized network activity across the interface. The development of the human RHT assembloid represents a significant leap forward in chronobiology. The “gold standard” for circadian models—self-sustained gene expression oscillations—was demonstrated using a *PER2::Luciferase* reporter, showing robust 20– 30 hour rhythms. This validates the hypothalamic component as a functional “clock in a dish”. This platform provides a readout to screen drugs or test light-pulse effects on circadian phase, directly modeling jet lag or phase-shifting. Overall, this model offers a high-fidelity system for investigating human-specific chronobiological mechanisms *in vitro*.

**Graphical Abstract:** 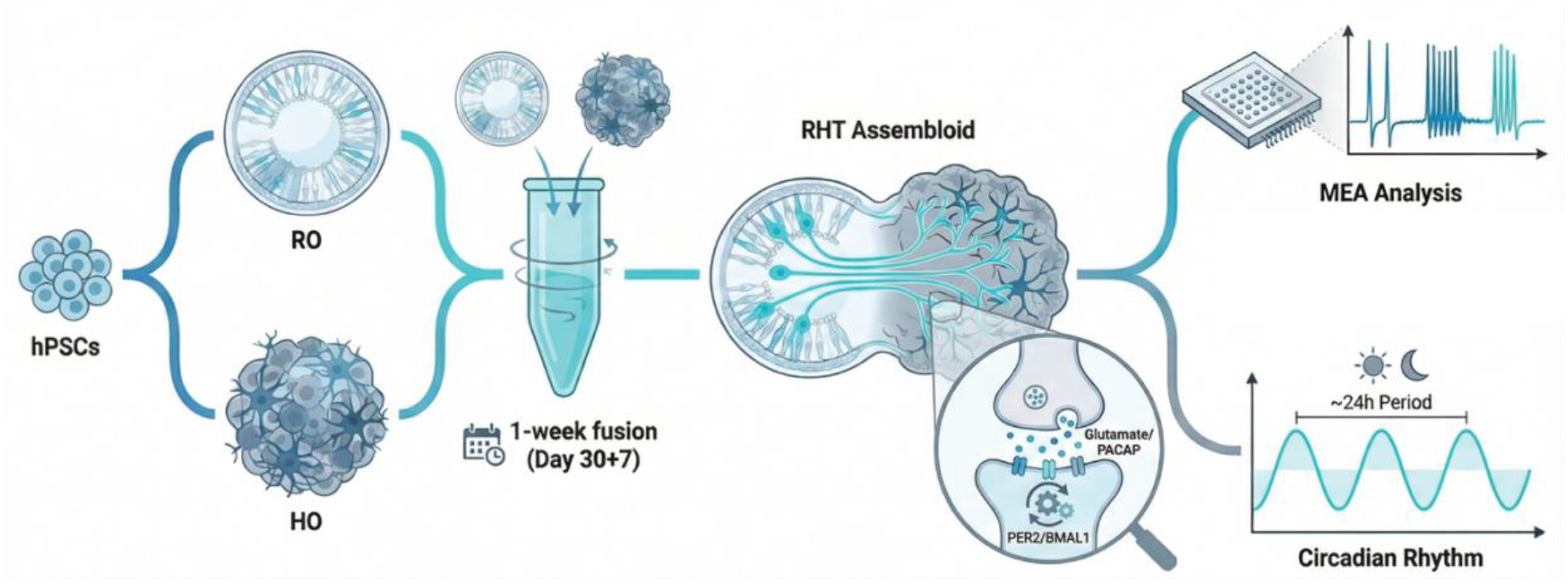

## Main

The retinohypothalamic tract is a pathway that extends directly from the retina to the suprachiasmatic nucleus (SCN) of the hypothalamus [1-3], and its primary function is to transmit photic information to the brain and play a role in non-image-forming (NIF) visual functions [2, 3]. Circadian rhythms are intrinsic biological oscillations that occur with a period of approximately 24 hours, coinciding with the Earth’s rotation, and are regulated by the master clock in the hypothalamic SCN [4, 5]. RHT functions as a dedicated input path that keeps the internal clock synchronized with the external light-dark cycle [6]. Traditional animal models such as mice face a significant limitation in studying human circadian disorders due to their nocturnal nature [7]. Similarly, clock gene regulation and circadian activity patterns differ between rodent models and day-active humans. Models derived from human induced pluripotent stem cells (iPSCs) overcome these drawbacks because they are derived from humans and the generated organoids incorporate circadian rhythms [8]. Organoids are three-dimensional structures that can be generated from pluripotent or adult stem cells that mimic human tissues cellularly and functionally [9-13]. Human iPSCs can be used to generate various neural organoids, including retinal organoids [14, 15], the brain [16-18], and sub-brain regions [19-22]. Retinal organoids (RO) produce mature photoreceptors that elicit strong, wavelength-specific light responses comparable to primate foveal cones [23]. Hypothalamus organoids (HO) also contain many of the cells that make up the hypothalamus [19, 20]. Co-culturing organoids yields more complex and functional assembloids [24-27]. It has been observed that retinal ganglion cells, which tend to decrease during maturation in RO, are preserved in greater numbers when combined with cortical and thalamic organoids to form assembloids [25]. Therefore, an assembloid model using RO and HO has the potential to be used for in vitro modeling of the human RHT. In this study, the retinohypothalamic tract assembloid model was created using ROs and HOs developed using human-derived PSCs. The RHT assembloid model recapitulates the human retina and hypothalamus with high similarity and has been characterized functionally The RHT model has a high degree of similarity to human physiology, both in terms of cellular content and electrophysiology. The model also possesses a circadian rhythm and can be used in circadian rhythm studies to overcome the disadvantages of traditional models. To our knowledge, this work establishes the first functional hiPSC-derived assembloid model of the human retinohypothalamic tract, providing a high-fidelity platform to investigate circadian photoentrainment and bridge the long-standing translational gap between rodent models and human physiology.

## Results

### Generation and Characterization of hiPSC-derived Retinal and Hypothalamus Organoids

To investigate the functional connectivity of the human retinohypothalamic circuitry, we first established a high-fidelity differentiation platform to generate human pluripotent stem cell hPSC-derived ROs and HOs. The retinohypothalamic tract is the primary pathway for circadian photoentrainment, relaying photic information from ipRGCs to the SCN of the hypothalamus (Fig. 1A). Using a temporal differentiation framework, we generated ROs and HOs over a 250-day period using stage-specific induction and maturation protocols (Fig. 1B). Morphological assessments showed that ciliary extensions of photoreceptors were clearly visible in ROs during the maturation phase (Day 200) (Figure 1C). We identified a critical milestone in photoreceptor maturation as the formation of a distinct ‘brush border’ at the apical margin (arrows). These hair-like protrusions correspond to nascent Outer Segments (OS), the specialized organelles housing the phototransduction machinery. The emergence of these structures indicates that the photoreceptors are not only structurally positioned but have developed the ultrastructural capacity for photon capture (Fig. 1C). In parallel, characterization of the hypothalamic lineage provided a clear sign of maturation; the neuroectodermal region exhibited progressive regression from Day 30 through Day 250 (Fig. 1D). This morphological refinement indicates a successful transition from initial progenitor expansion to specialized neuronal differentiation. To validate the biological fidelity and spatial organization of the ROs, we performed longitudinal immunofluorescence analysis for recoverin, a mature photoreceptor marker. Recoverin is a canonical marker for both rod and cone photoreceptors and its expression indicates the recovery phase of phototransduction. By Day 200, the organoid achieves a critical architectural milestone: the establishment of a discrete, peripherally located photoreceptor layer (Fig. 1E). This specific organization maximizes the tissue’s ability to capture light. Since these recoverin-positive cells provide the extrinsic light signals to the inner retinal circuitry, including RHT, their proper spatial positioning within the RHT assembloid model and maturation are essential for modeling accurate circadian rhythm responses *in vitro*.

**Figure 1.**
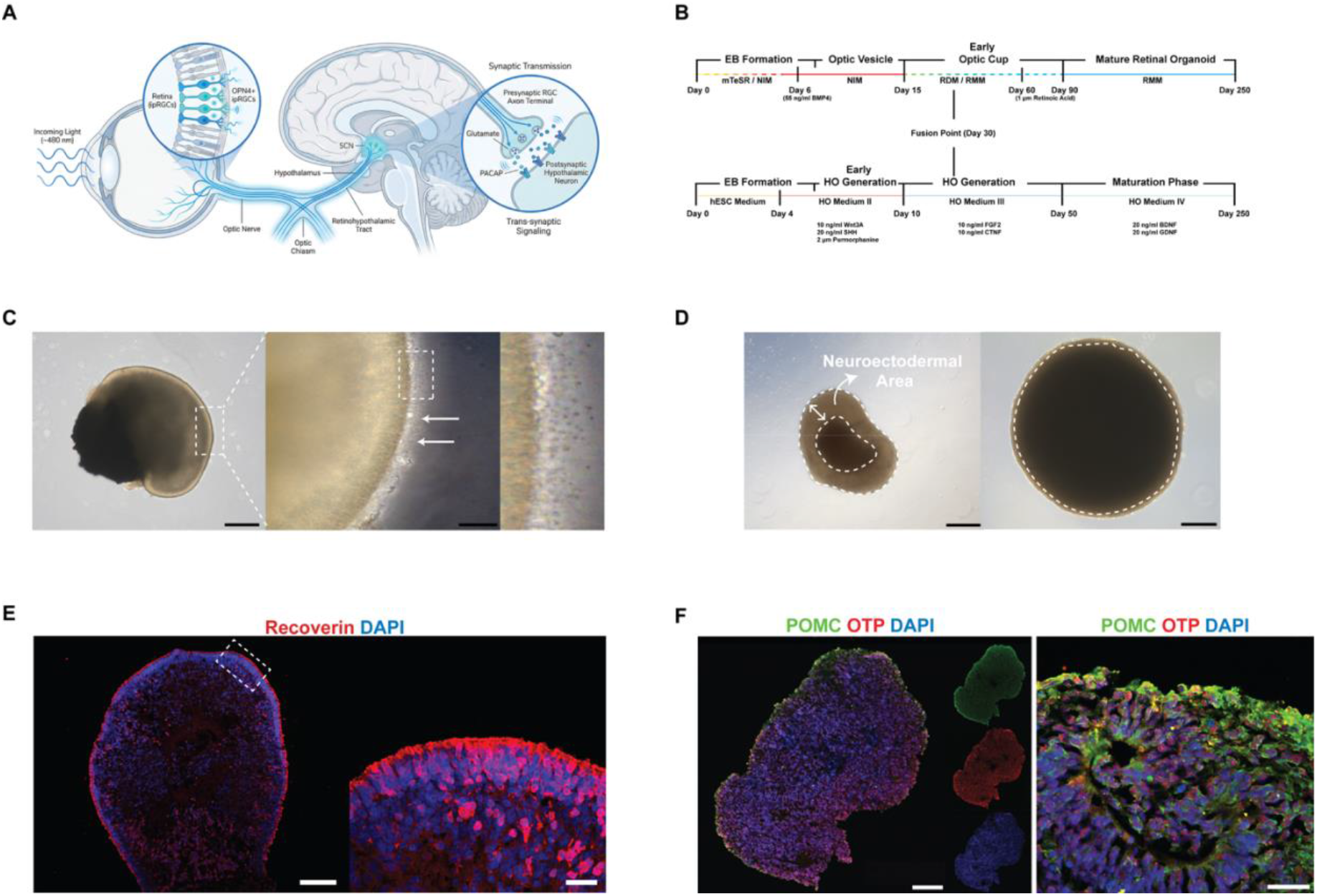
Generation and characterization of hPSC-derived retinal and hypothalamus organoids. **(A)** Schematic illustration of the human retinohypothalamic tract. This pathway mediates circadian photoentrainment by transmitting light signals from the retina to the suprachiasmatic nucleus in the hypothalamus. **(B)** Experimental workflows and timelines for the differentiation of RO and HO from hPSCs. **(C)** Brightfield microscopy of RO maturation. High-magnification insets at Day 200 reveal the formation of a distinct apical ‘brush border’ (arrows), corresponding to nascent photoreceptor outer segments (OS). **(D)** Morphological progression of HOs from Day 30 to Day 250. Arrows denote the progressive regression of the neuroectodermal region, indicating the successful transition from progenitor expansion to neuronal differentiation. **(E)** Immunofluorescence analysis of ROs stained for the mature photoreceptor marker recoverin (red). **(F)** Molecular validation of HO identity and diversity at Day 250. Positive staining for the hypothalamic progenitor marker OTP (red) and peptidergic neuron marker POMC (green) confirms the presence of specialized hypothalamic cell populations. Nuclei are counterstained with DAPI (blue). Scale bars, 400 µm (**c** (left), **d**), 200 µm (**f** (left)), 100 µm (**c** (right), **e** (left)), 20 µm (**f** (right)), 10 µm (**e** (right)).

Furthermore, the cellular identity and diversity of HOs were validated through the expression of the peptidergic neuron marker (POMC) and the hypothalamic progenitor marker (OTP) (Figure 1F). These findings demonstrate that the hPSC-derived ROs and HOs successfully recapitulate the morphological and molecular hallmarks of human tissue development, providing a robust foundation for the RHT assembloid model.

### RHT Assembloid Generation and Characterization

To investigate the structural and functional integration of the human retinohypothalamic circuitry, we generated assembloids by fusing Day 30 ROs and HOs using an optimised co-culture protocol. The assembly process followed a predefined timeline, transitioning from initial fusion to long-term maturation in specialized assembloid media (Fig. 2A, B). By Days 190 after fusion (D.A.F. 190), brightfield microscopy confirmed the fusion of the two distinct orgaoids while maintaining clear morphological boundaries (Fig. 2C). Longitudinal morphological assessment of assembloids generated from the H9 human embryonic stem cell line confirmed that the initial physical contact established at Day 30 stabilized into a well-defined fusion interface by D.A.F. 7. This structural integrity was robustly maintained throughout the intermediate maturation phase at D.A.F. 60, with the distinct tissue components remaining tightly fused without loss of tissue-to-tissue contact (Fig. 2D). We further validated the structural integrity of the assembloid through regional marker analysis, where recoverin positive photoreceptors were strictly confined to the retinal side (Fig. 2F). Recoverin expression was exclusively localized to the retinal component, demonstrating that the fused tissues maintain their distinct cellular identities without aberrant intermingling throughout the maturation process. (Fig. 2F). This architectural stratification is a critical prerequisite for establishing directed signal transmission from the sensory retinal layer to the hypothalamic core (Fig. 2F).

**Figure 2.**
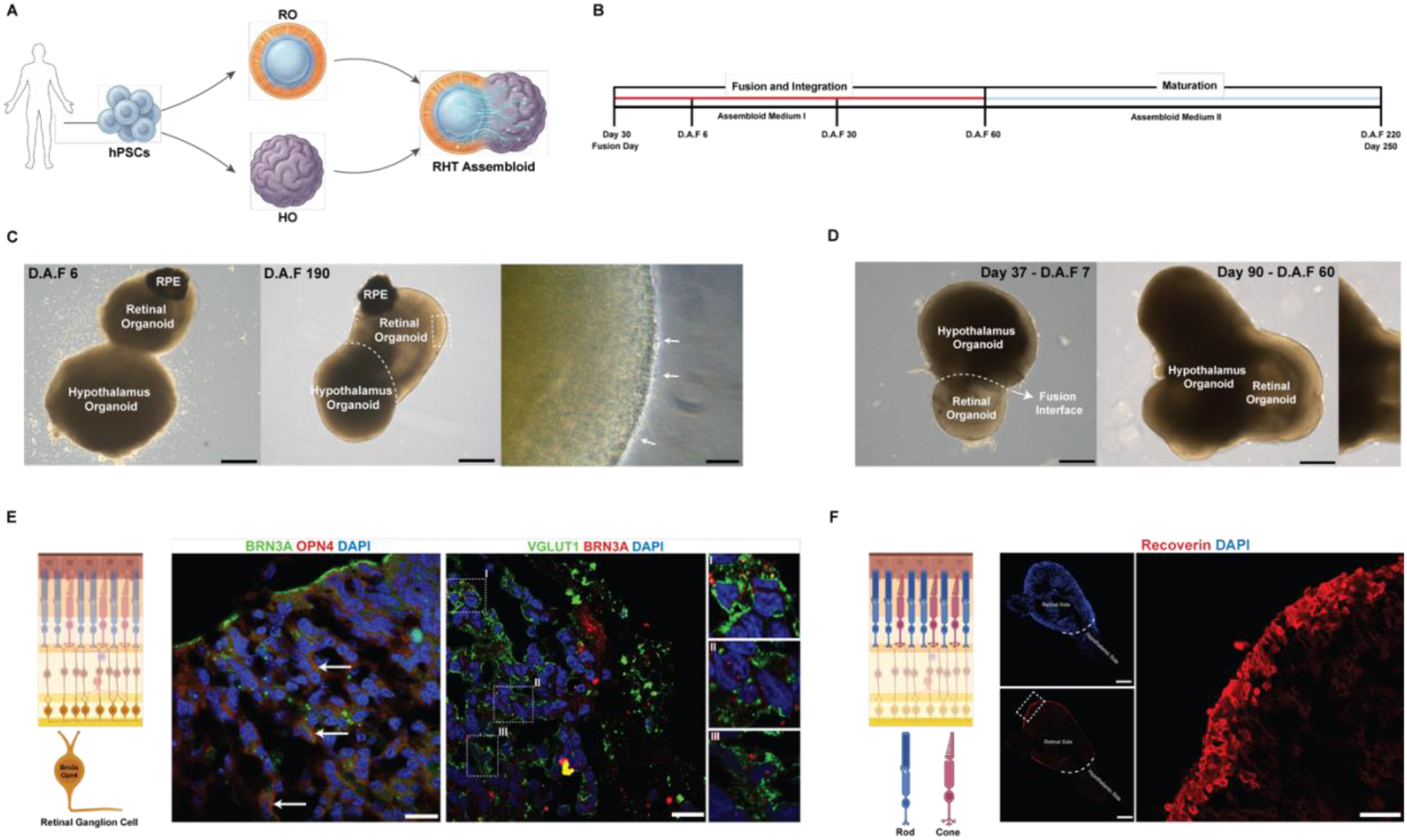
Generation and characterization of the human RHT assembloid model. **(A)** Schematic of RHT assembloid generation. Overview of the experimental workflow starting from hPSCs to the differentiation of RO and HO. The inset shows a magnified view of synaptic integration between retinal ganglion cell RGC axons and hypothalamic neurons. **(B)** Experimental timeline. Sequential stages of fusion and integration (Day 30 to D.A.F. 60) followed by a long-term maturation phase in specialized assembloid media, extending to D.A.F. 220 (Day 250). **(C)** Morphological assessment of fusion. Representative brightfield images of the assembloid at D.A.F. 6 showing initial tissue contact and at D.A.F. 190 showing advanced structural integration and the presence of pigmented retinal pigment epithelium and peripheral photoreceptor layers (arrows). **(D)** Longitudinal brightfield microscopy of H9 hESC-derived assembloid maturation. Representative images at D.A.F. 7 (total Day 37) and D.A.F. 60 (total Day 90) demonstrate the progressive stabilization of the fusion interface and the maintenance of structural continuity between the RO and HO. **(E)** Characterization of ipRGCs and excitatory synapses at D.A.F. 190. (Left) Immunofluorescence staining for BRN3A (green) and Melanopsin (OPN4, red) confirms the presence and long-term survival of intrinsically photosensitive retinal ganglion cells (ipRGCs), essential for circadian entrainment. (Right) Expression of the excitatory marker VGLUT1 (green) in BRN3A+ RGCs. The punctate VGLUT1 staining pattern suggests the formation of mature glutamatergic presynaptic terminals ready for signal transmission. Nuclei are counterstained with DAPI (blue). **(F)** Maintenance of tissue stratification and regional identity at D.A.F. 190. Immunofluorescence analysis showing that recoverin-positive photoreceptors (red) are strictly confined to the retinal side of the assembloid, with no expression observed on the hypothalamic side. This confirms that fused tissues maintain their distinct cellular identities and structural integrity without aberrant intermingling. Nuclei are counterstained with DAPI (blue). Scale bars, 400 µm (**c** (left, mid), **d**), 100 µm (**f** (left, mid)), 40 µm (**c**(right)), 20 µm (**f** (right)), 10 µm (**e**).

### Identification of ipRGCs and Synaptic Maturation at the Assembloid Interface

The establishment of a functional retinohypothalamic tract *in vitro* necessitates the maturation of ipRGCs and their associated excitatory machinery. Analysis at D.A.F. 190 confirmed the survival and maturation of these specific neuronal populations within the integrated circuitry. We identified a robust population of retinal ganglion cells through positive staining for BRN3A (Fig. 2E). Crucially, the presence of melanopsin (OPN4) expression within these cells confirmed the existence of ipRGCs [28], which are essential cellular components for non-image forming visual functions, specifically circadian entrainment (Fig. 2E). To assess the readiness for signal transmission, we examined the expression of the vesicular glutamate transporter VGLUT1, which revealed an excitatory glutamatergic profile (Fig. 2E). The characteristic punctate staining pattern of VGLUT1 surrounding the BRN3A+ RGC nuclei suggests the formation of mature presynaptic terminals, indicating that the assembloid model has developed the necessary synaptic infrastructure for functional excitatory output from the retina to hypothalamic neurons (Fig. 2E).

### Electrophysiological Maturation and Functional Synaptic Architecture of Hypothalamus organoids and RHT assembloids

To determine the functional maturation and network-level activity of the generated tissues, we combined high-resolution immunohistochemistry with MEA recordings. Initial assessment of Day 30 HOs through immunostaining for synaptophysin revealed a highly organized spatial distribution of synaptic machinery (Fig. 3A). While the surrounding neuropil exhibited dense accumulations of synaptic vesicles, the apical neural rosettes— which serve as progenitor zones—remained largely devoid of synaptic markers (Fig. 3A). This distinct pattern mimics the *in vivo* developmental separation between the proliferative ventricular zone and the differentiating mantle zone, confirming the structural fidelity of our HO model. Longitudinal characterization of the synaptic protein profile further revealed a progressive upregulation of key molecular components within the HO circuitry. We observed a significant increase in CTBP2 intensity from Day 30 to Day 90, confirming the robust recruitment of essential synaptic machinery during organoid maturation (Fig. 3B). Interestingly, the recruitment of Bassoon appeared temporally delayed relative to CTBP2, suggesting that the consolidation of the active zone scaffold is a subsequent maturation step that follows the initial accumulation of ribbon components at the presynaptic terminal (Fig. 3B).

**Figure 3.**
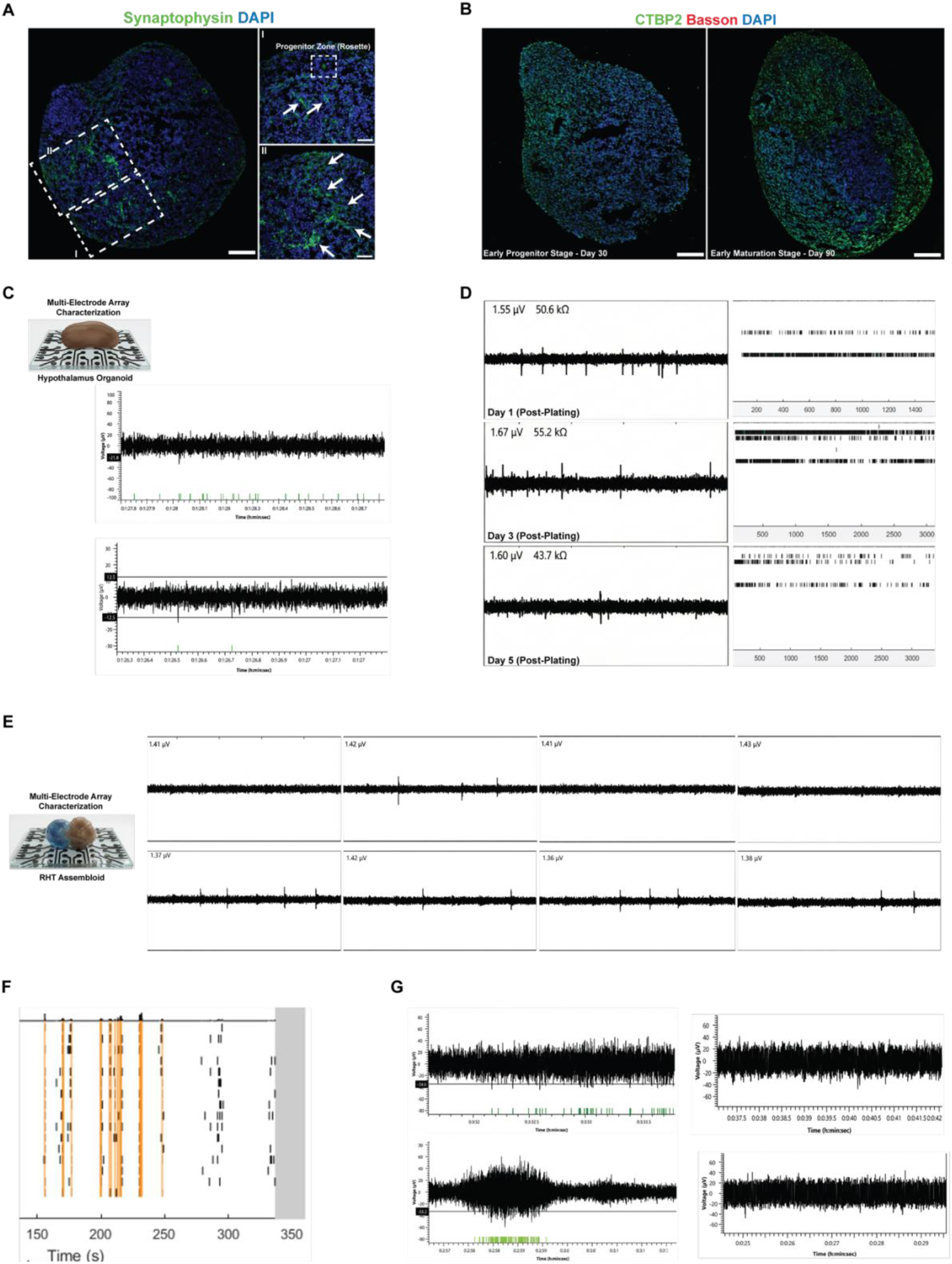
Electrophysiological maturation and synaptic architecture of HOs and RHT assembloids. **(A)** Synaptophysin expression in Day 30 HOs showing spatial separation between synaptic neuropil and progenitor rosettes. **(B)** Longitudinal IF analysis from Day 30 to Day 90 showing progressive CTBP2 upregulation and the subsequent recruitment of Bassoon to the active zone scaffold. **(C-D)** MEA recordings of spontaneous activity in HOs at Day 75 and Day 180, showing increased spike density and burst frequency. **(E-G)** Functional validation of RHT assembloids at D.A.F. 45 via MEA, including raster plots (F) and localized trace analysis (G) demonstrating integrated network activity. Scale bars, 100 µm (**a** (left), **b**), 50 µm (**a** (mid, right)).

### Progression of Spontaneous Activity and Network Integration

The electrophysiological development of the HOs was tracked longitudinally using MEA analysis to evaluate spontaneous firing patterns. At Day 75, HOs exhibited baseline spontaneous activity characterized by isolated spikes (Fig. 3C). By Day 180, a significant increase in spike frequency and burst complexity was observed, indicating successful long-term neuronal maturation and the establishment of more complex local circuits (Fig. 3D). The functional integration of the RHT assembloid was evaluated at 45 days after fusion (D.A.F. 45; corresponding to total Day 75). MEA recordings from these assembloids demonstrated robust and continuous spontaneous activity across the fused interface (Fig. 3E). Raster plot analysis revealed synchronized firing patterns and organized burst activity, suggesting the formation of an integrated neural network between the retinal and hypothalamic tissues (Fig. 3F). Detailed localized recordings confirmed that the physical assembly of ROs and HOs resulted in a functionally active circuit capable of maintaining stable electrophysiological output, a critical prerequisite for investigating light-induced signal transmission (Fig. 3G).

### Molecular Characterization of the SCN Master Clock and Autonomous Circadian Oscillations

To determine if the RHT assembloid model possesses a functional biological clock, we first investigated the expression of the core transcriptional-translational feedback loop (TTFL) within the hypothalamic SCN region. The molecular clockwork, driven by the reciprocal regulation of CLOCK and BMAL1 activators and PER/CRY repressors, serves as the fundamental pacemaker for ∼24-hour rhythms (Fig. 4A). To enable the real-time monitoring of these molecular oscillations, we established a luciferase-reporter system using a validated iPSC line. To ensure the reliability of this monitoring platform, we examined the luciferase activity across multiple cell passages; the results demonstrated a stable and consistent light unit output, confirming that the reporter system maintains its technical integrity and sensitivity for longitudinal circadian recording (Fig. 4B). With the monitoring system validated, we next confirmed the establishment of SCN identity within the HOs through the expression of Vasoactive Intestinal Peptide (VIP), a hallmark marker of the master pacemaker. Longitudinal analysis revealed that VIP expression became significantly more organized and robust in the mature neuropil by Day 250 compared to early stages (Fig. 4C). Further characterization confirmed the presence of the complete core clock machinery, with distinct nuclear localization of BMAL1, CLOCK, CRY1, and PER1 within the hypothalamic neurons (Fig. 4D). Finally, the functional autonomy of the clock was evaluated via real-time bioluminescence recording. RHT assembloids exhibited robust, self-sustained circadian oscillations in PER2 activity, with Cosinor analysis confirming a stable rhythm within the biological 20–30 hour range (Fig. 4E).

**Figure 4.**
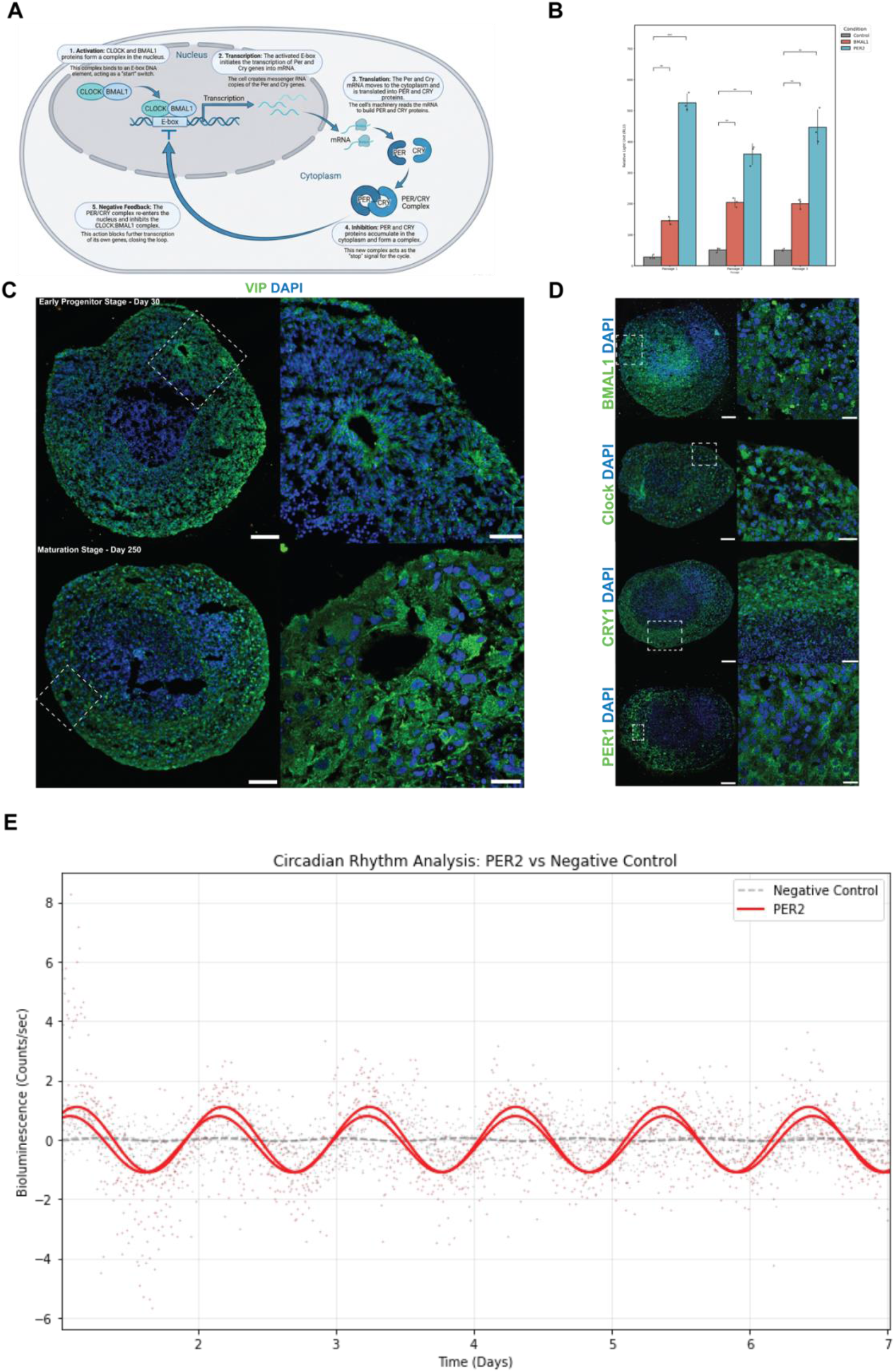
Molecular architecture and functional validation of the autonomous circadian clock. **(A)** Schematic of the core circadian transcriptional-translational feedback loop (TTFL). **(B)** Technical validation of the reporter system. Stable relative light unit (RLU) output across consecutive passages of the iPSC ensures the reliability and consistency of real-time monitoring. **(C)** Immunofluorescence staining for VIP at Day 30 and Day 250, showing the maturation of SCN-like peptidergic identity. **(D)** Representative images showing nuclear localization of core clock proteins: BMAL1, CLOCK, CRY1, and PER1. **(E)** Real-time circadian rhythm analysis. Baseline-subtracted bioluminescence data (red dots) for PER2 show robust oscillations with a stable Cosinor fit (solid red line) compared to the negative control (dashed line). Scale bars, 100 µm (**c** (top and bottom left side), **d** (all left side)), 50 µm (**c** (top right side), **d** (third line right side)), 20 µm (**d** (second line right side)), 10 µm (**c** (bottom right side), **d** (first and last line right side)).

## Discussion

Circadian rhythms are fundamental to human health, as their desynchronization is linked to increased risks of metabolic, cardiovascular, and neurodegenerative disorders. Central to this regulation is the retinohypothalamic tract, a direct neural pathway that transmits photic signals from specialized retinal photoreceptors to the SCN to align internal physiology with the environmental light-dark cycle (Fig. 1A). Historically, our understanding of the retinohypothalamic circuitry has relied heavily on model organisms such as nocturnal rodents. However, these models exhibit a significant translational gap due to fundamental differences in chronotypes; humans are diurnal, meaning the phase relationships of core clock gene expression and physiological responses to light are often inverted compared to nocturnal species [8, 29, 30]. Furthermore, anatomical and molecular differences in human ipRGC subtypes and their projection patterns to the SCN suggest that rodent-based entrainment mechanisms do not fully recapitulate human photic biology [7]. Traditional 2D cell cultures, while useful for studying cell-autonomous molecular clocks, fail to recapitulate the complex 3D tissue architecture and the heterogeneous cellular niches required for systemic circadian synchronization. Rodent models have been invaluable, but their nocturnal nature means that the “output” of their clock (e.g., activity, melatonin) is anti-phase to humans. Furthermore, human retina possess unique ipRGC subtypes and distinct spectral sensitivities [31]. While recent advances in organoid and assembloid technologies have provided a “new window” into human brain development, existing models have primarily focused on the intrinsic rhythmicity of individual regions. For instance, circadian oscillations have been independently demonstrated in human cerebral organoids [32] and ROs [33]; however, these isolated systems represent only partial complexity and lack the integrated functional circuitry of the RHT. In this study, by assembling the direct elements of human circadian regulation into a functional RHT assembloid, we have established a high-fidelity platform that closely mimics human physiology and provides a robust model for investigating chronobiological mechanisms *in vitro*. The development of the human RHT assembloid model represents a significant leap forward in the field of chronobiology, addressing the long-standing translational gap between rodent research and human physiology. By utilizing human iPSCs, this model captures the human-specific genomic and proteomic landscape of the RHT. For instance, the specific isoforms of Melanopsin or the precise stoichiometry of synaptic proteins in human ribbon synapses can now be studied in a native context.

The emergence of a distinct apical brush border at Day 200 (Fig. 1C) represents a critical structural hallmark of advanced retinal maturation. This ultrastructural specialization, corresponding to nascent outer segments (OS), is consistent with the developmental milestones previously identified in high-fidelity RO models [34]. While classical rod and cone photoreceptors provide redundant signals to the circadian system alongside ipRGCs [30], their well-defined spatial organization in our model underscores the establishment of a sensory-competent retinal architecture required for accurate photoentrainment. The formation of a distinct peripheral layer of Recoverin-positive photoreceptors by Day 200 (Fig. 1E) represents a critical stage in the establishment of mature retinal architecture [35]. This organization mirrors the formation of the Outer Nuclear Layer (ONL) in the human retina, enabling the photoreceptor cell bodies to separate from the inner neuroepithelium and stratify in an organized manner. This observed laminated structure is in complete agreement with high-fidelity organoid models that mimic human retinal development and structural maturation in long-term cultures [34, 36]. Morphological development observed in our HO demonstrated a significant regression of neuroectodermal areas between Day 30 and Day 250, giving way to mature neuronal tissues (Fig. 1D). This morphological refinement is a strong indication that hypothalamic progenitor cells have transitioned from the proliferative phase to differentiate into specific neuronal subtypes. In the literature, the time-bound narrowing of ventricular zone-like neuroepithelial structures in human iPSC-derived neural models is considered a successful maturation criterion [16, 17, 37]. Characterization of HOs in Day 250 revealed the co-occurrence of the hypothalamic progenitor marker OTP and the mature peptidergic neuron marker POMC, exhibiting an organized distribution (Fig. 1F). The presence of POMC+ neurons and OTP+ progenitor regions in HOs successfully mimics the transition from progenitor expansion to neuronal enrichment in human hypothalamic development [19, 21]. Also, co-expression or distinct localization of these markers (Figure 1F) validates the cellular diversity of the HOs, suggesting they contain the necessary nuclei (e.g., SCN-like) to form a functional hypothalamic circuit.

The RHT assembloid model was created (Fig. 2 A-B) in such a way that RGCs extend into the hypothalamic tissue, as in human RHT. In isolated ROs, RGCs differentiate early (around Day 40–50) but typically undergo apoptosis between Day 90 and 120 [14, 38]. At D.A.F 160 (Day 190), BRN3A+ cells were observed to be conserved in the RHT assembloid model (Figure 2E). This situation has been shown to contribute to the long-term survival of the assembloid model of the RGC [25], and this survival is significantly beyond the typical window of loss in isolated cultures, providing strong evidence that the HO provides the necessary trophic support to rescue the RGC population. This study identified ipRGCs, a specific subtype of RGCs that express melanopsin [28] in the RHT assembloid model (Fig. 2E). ipRGCs are the specific conduit for non-image-forming vision, including circadian photoentrainment. Unlike rods and cones, which hyperpolarize in response to light, ipRGCs depolarize directly via an opsin-based phototransduction cascade. The detection of melanopsin protein at Day 190 confirms that the human RHT model contains the specific cellular hardware required for circadian light detection. The presence of these cells suggests that the assembloid could be used to study the specific effects of blue light (approx. 480 nm, the peak sensitivity of melanopsin) on hypothalamic physiology [39]. The RHT is an excitatory glutamatergic pathway. VGLUT1, considered a marker of excitatory presynaptic terminals responsible for glutamate loading into synaptic vesicles, is expressed in the RHT assembloid model close to BRN3A+ RGCs (Fig. 2E). This implies that the model can generate functional oscillation zones, which is consistent with recent findings in thalamocortical assembloids [40]. Following organoid fusion, recoverin+ cells remained confined to the retinal region in assembloids and did not undergo abnormal migration to the hypothalamus or vice versa (Fig. 2F). This successfully mimics the separation of the retina from the hypothalamus in vivo, implying communication via optic nerve-like axonal pathways rather than a complex structure, and allowing for precise transmission of directional signals [41].

To prove functional competence, the assembloids were functionally validated by immunochemical verification and synaptic protein analysis, as well as by MEA (Fig. 3). CTBP2, also known as RIBEYE, is a marker of the synaptic nucleus, while Basson is the active site protein that binds the strip to the presynaptic membrane. Synaptophysin is an integral synaptic vesicle protein. Immunochemical verification of these provides preliminary characterization of the electrophysiological activity of the organoid and indicates its preservation in long-term culture (Fig. 3A-B). Furthermore, these indicators show that the organoids have achieved the ability to maintain high tonic release rates necessary for visual signaling [42]. In HOs, sparse spikes obtained by MEA on day 75 evolved into high-frequency, complex bursts on day 180, a classic indicator of neuronal network maturation (Fig. 3C-D). Synchronized activities (Fig. 3E-G) detected in RHT assembloids and raster plots (Fig. 3F) show a vertical alignment of firing events across multiple electrodes. This “network bursting” implies that the neurons are not firing independently but are connected in a coordinated circuit. This synchronization potentially demonstrates that spontaneously generated retinal waves can propagate to other regions. For RHT, this connection is a functional prerequisite for synchronization, and spontaneous retinal activity, even in the absence of light, plays a crucial role in the developmental connections of the visual system and the SCN. Assembloid successfully models this metanetwork connection [43].

The mammalian circadian clock is regulated by the Transcriptional-Translational Feedback Loop via the Bmal1, Clock, Period (PER), and Cryptochrome (CRY) genes. SCN neurons synchronize with specific neuropeptides. Vasoactive Intestinal Peptide is one of the most critical of these, and VIP-expressing neurons in the ventral SCN receive direct RHT input and transmit this information to the dorsal SCN sheath. Our HOs show strong VIP expression, becoming more organized by day 250 (Fig. 4C). Without VIP signaling, individual SCN neurons function like independent clocks with different periods, leading to a damped, “flat” tissue-level rhythm. Therefore, this peptide is essential for a model in which the circadian rhythm can be studied. Our HOs also confirm the expression and, critically, nuclear localization of BMAL1, CLOCK, CRY1, and PER1 (Fig. 4D). Translocation of PER/CRY complexes to the nucleus is the critical step that inhibits CLOCK/BMAL1 activity and closes the feedback loop. Visualization of these proteins in the nucleus confirms that the “repression phase” of the clock is active. This indicates that the organoids not only express clock genes but are also subject to the dynamic subcellular trafficking necessary for oscillation [8].

The functional “gold standard” for any circadian model is the demonstration of self-sustained oscillations in gene expression. Using a *PER2::Luciferase* reporter line, the study demonstrates robust bioluminescence cycles with a period of 20–30 hours (Figure 4E). The “Cosinor fit” confirms that the rhythm is statistically significant and stable over multiple days. This autonomous oscillation is a defining feature of the SCN. Unlike peripheral tissues, which dampen quickly in isolation, the SCN (and this organoid model) can sustain high-amplitude rhythms indefinitely due to its internal coupling. This finding validates the HO component as a functional “clock in a dish.” It provides a readout (luminescence) that can be used to screen drugs or test the effects of light pulses (delivered via the attached retinal organoid) on circadian phase, directly modeling the mechanism of jet lag or phase-shifting [32].

## Methods

### Pluripotent Stem Cell Culture

hiPSCs [44] and hESCs were cultured in 1% embryonic stem cell (ESC) qualified Matrigel (Corning) 6-well plates using mTeSR1 (STEMCELL Technologies) medium at 37 °C, 5% CO2, in a humidified incubator. The medium was changed daily until cell density reached 80%. Cells were passaged 1:10 using ReLeSR (STEMCELL Technologies).

### Lentiviral packaging and transduction

HEK293T cells were seeded into the wells of a 6-well plate to reach approximately 80% confluence after 24 hours and were co-transfected with a target vector (pLV7-Bsd-p(Bmal1)-KH-dLuc or pLV7-Bsd-p(Per2)-KB-dLuc) and two helper plasmids, psPAX2 (Addgene plasmid #12260) and pMD2.G (Addgene plasmid #12259), at a ratio of 4:2:1, using a total DNA amount of 2.5 μg. Transfection reagent, Polyethylenimine (PEI; 23966, Polysciences), was used at a 1:5 DNA-to-PEI ratio. Cell media were collected 48 h after transfection and filtered through a 0.45 μm SFCA filter (431220, Corning). IPS cells were seeded at ∼50–60% confluency and transduced for 48 h with medium containing viral particles diluted in complete growth medium supplemented with polybrene (H9268, Sigma Aldrich) at a final concentration of 8 μg/ml. Infected cells were selected with 1 μg/ml puromycin (ant-pr-1, InvivoGen) for 72 hours.

### Retinal Organoid Generation

Retinal organoid generation protocol developed by Fligor et al. was used to generate ROs [14]. The stages of generation ROs are divided into three steps: neural induction process, induction to a primitive retinal fate and generation of organoids from primitive retinal cells. First, 5 wells of a six well plate that were 80% confluent were cultured in a 75 ml flask for 6 days for retinal differentiation. A gradual transition from the iPSC medium mTeSR to neural induction medium (NIM) was made for 6 days to give neural characteristics to the cell aggregates. On the 6th day, the retinal differentiation process was initiated by adding BMP4 (55 ng/ml) to NIM. To induce primitive retinal fate in iPSC aggregates, cell aggregates are transferred from suspension culture to 2D culture. BMP4-containing medium is gradually replaced with BMP4-free medium until day 16. On day 16, cell aggregates are transferred back to 3D culture and retinal differentiation medium (RDM) is added. On day 35, taurine is added to RDM to support photoreceptor cell formation. Retinoic acid is added between day 60 and day 90 to maintain and maturate photoreceptor cells. From day 90 onward, retinal maturation medium (RMM) was applied to promote further differentiation and maturation of ROs.

### Hypothalamus Organoid Generation

Hypothalamus organoids were generated by modifying the protocol developed by Qian et al [19]. The process of generating HOs was completed in three stages: embryoid body (EB) formation, early HO formation, and maturation. EBs were cultured in u-bottom ultralow attachment 96-well plates with approximately 10,000 cells per well. PSCs were cultured with hESC medium every other day for the first four days [17]. HO2 medium was applied between days 5 and 10. From day 10 onward, EBs with a diameter of 500–60 µm were transferred to ultralow attachment 24-well plates and started receiving HO3 medium. On day 50, HOs were transferred to ultralow attachment 6-well plates and switched to HO4 medium.

### RHT Assembloid Generation

We used the method developed by Sloan et al. to generate assembloids from retinal and hypothalamus organoids [24]. We incubated Day 30 organoids in 1.5 ml microcentrifuge tubes. We incubated for 1 week, changing the medium every 2 days. Upon completion of fusion, we transferred the assembloids to ultralow attachment 6-well plates.

### Immunohistochemistry

Organoids were fixed with 4% paraformaldehyde (Sigma-Aldrich) for 24 hours at 4 °C and then embedded in cryomatrix (OCT, Fisher Healthcare). Cryosections were taken with Leica CM1950 (Leica Biosystems, Germany). Sections were permeabilized with permeabilization buffer (0.3% Triton P7949) (v/v), 1% Bovine Serum Albumin (BSA; Sigma-Aldrich, Cat no. A2153) (w/v) in PBS) were added to the relevant samples and incubated overnight at 4 °C. Cells were stained with relevant secondary antibodies for 2 h at room temperature. Nuclei were stained with 4,6-diamidino-2-phenylindole. (0.5 μg/mL) (DAPI; Neofroxx, Cat no. 1322). Images of the samples were captured with a confocal microscope (Zeiss LSM880).

### Organoid culture on MEAs

To record extracellular activity from organoids, microelectrode arrays (MEAs; electrode diameter: 30 μm; inter-electrode spacing: 100 μm; MEA256 100/30 iR-ITO, MultiChannel Systems) and an MEA2100 amplifier were used for data acquisition. Prior to organoid culture, MEA surfaces were sterilized by rinsing with 70% ethanol for 5 min, followed by ultraviolet (UV) exposure for 30 min. Sterilized MEAs were coated overnight with poly-L-lysine (PLL; 0.1 mg/mL; Sigma-Aldrich, MO, USA) to promote cell adhesion and then washed with nanopure water to remove excess PLL. Retinal or hypothalamic organoids, as well as RTH assembloids, were placed onto PLL-coated MEA chips and maintained in culture. Electrophysiological recordings were performed after 3 days of culture after complete attachment to the surface. Extracellular neuronal activity was acquired using a 256-channel amplifier and recorded with MultiChannel Experimenter software (MultiChannel Systems).

Also, HOs, and assembloids were plated per well in 48-well MEA plates (Axion Biosystems, USA). Each well contains 16 low-impedance (0.04 MΩ) platinum microelectrodes. The plate was previously coated with 100 μg/mL poly-L-ornithine and 10 μg/mL laminin. Cells were fed once in three days, and electrical activities were monitored for two weeks. Electrical activities were recorded using a Maestro MEA system and AxIS Software Spontaneous Neural Configuration (Axion Biosystems) with a customized script for band-pass filter (0.1-Hz and 5-kHz cutoff frequencies). Spikes were detected with AxIS software using an adaptive threshold crossing set to 6 times the standard deviation of the estimated noise for channel. The plate was first allowed to rest for 3 min in the Maestro device, and then data were recorded.

### MEA data analysis

Raw extracellular recordings were acquired from a 256-channel microelectrode array using the MEA2100 system (Multi Channel Systems) at an effective sampling rate of 10 kHz. Samples were allowed to equilibrate on the MEA for at least 20 min prior to recording. Raw electrophysiological signals were imported from HDF5 files, band-pass filtered between 300 and 3000 Hz, and analyzed offline using MATLAB R2025b. Spike detection was performed using a threshold-based method defined as −4.5× the standard deviation of baseline noise. Electrodes exhibiting negligible activity (mean firing rate < 0.02 Hz or fewer than five detected spikes during the recording period) were excluded from further analysis. Spatial activity maps were generated by projecting electrode-wise mean firing rates onto the physical 16×16 layout of the MEA.

### Organoid-based Luminescence Assay and Circadian Rhythm Analysis

Organoid-based circadian rhythm experiments were conducted using size-matched organoids rather than single-cell cultures. For each experimental condition, two assembloids *BMAL1*-d*Luc*/*PER2*-d*Luc* of comparable size were selected to minimize variability associated with differences in organoid volume and cellular composition. Prior to luminescence recording, attachment-promoting components on the surface of tissue culture–treated polystyrene (TC-treated PS) plates were neutralized to enable free-floating conditions. Briefly, 35-mm culture plates were incubated with Anti-Adherence rising solution for 4 hours to reduce protein-mediated surface adhesion to prevent premature attachment and to support 3D aggregation, as demonstrated previously in acoustic assembly and culture systems.

Circadian synchronization was achieved by treating organoids with dexamethasone (DEX) at a final concentration of 0.1 μM in DMEM (1 ml per plate) [45, 46]. After a 2-hour incubation at 37 °C, the medium was removed and replaced with freshly prepared luminescence recording medium. The recording medium consisted of 10 g DMEM powder (Sigma, D-2902), 0.35 g sodium bicarbonate (Sigma, S5761), and 3.5 g D-(+)-glucose powder (Sigma, G7021), dissolved in distilled water. HEPES buffer (1 M; Gibco, Cat. No. 15140-122) was subsequently added to achieve a final concentration of 10 mM. Penicillin–streptomycin (Capricorn Scientific, PS-B) and fetal bovine serum (Gibco, Cat. No. A5256801) were added to final concentrations of 0.25% (v/v) and 5% (v/v), respectively. Non-essential amino acids were supplemented at a final concentration of 1% (v/v). The volume was adjusted to 1 L with distilled water, and the medium was sterilized using a 0.22 μm Millex MP filter unit. D-luciferin was freshly added immediately before use at a final concentration of 0.1 mM [45, 46]. The recording medium was added to the organoids at a volume of 2 ml per plate. Plates were sealed with lids coated with silicon grease to prevent evaporation and then placed into the LumiCycle system (Acrimetrics) for continuous monitoring of the circadian rhythm. Bioluminescence signals driven by the BMAL1 and PER2 promoters were recorded for 5–7 days. Data were collected every 10 minutes at 37 °C, with a 75-second integration time per measurement. The resulting data were analyzed using LumiCycle Analysis software (Acrimetrics) to determine circadian parameters, as previously described [46]. Thereby characterizing the circadian dynamics of PER2 promoter activity in organoid cultures.

## Acknowledgements

This work is supported by TÜBİTAK 124Z226 project. B.K. is fellow of TÜBİTAK 2211C and TÜBİTAK 2250 scholarship program. We gratefully acknowledge the Histology and Optical Imaging Core Facilities at the İzmir Biomedicine and Genome Center (IBG) for their expert support in tissue preparation and advanced confocal microscopy. Figures 2E and 2F were created with biorender.com. During the preparation of this article, the authors used Gemini and NotebookLM to improve the text flow and better present the information. The information obtained from Gemini was reviewed and corrected by the authors. They took full responsibility for the content of the published article. The authors express their gratitude to the Koç University Research Center for Translational Medicine (KUTTAM) for providing access to its services and facilities, supported by funding from the Republic of Turkey Ministry of Development. The content is solely the responsibility of the authors and does not necessarily represent the official views of the Ministry of Development. H.N.K. has financial support from the TÜBİTAK 2218 scholarship program.

## Author Contributions

B.K. and S.G. established the hypothesis and study design. The hypothalamus, retinal organoids, and assembloids were constructed and characterized by B.K. The generation of BMAL1 and PER2 Luciferase reporter lines from iPSCs was carried out by E.Ç and Ş.Ş. MEA recording and analyses were performed by B.D, H.N.K, Ö.F.Ö, S.N, H.B and B.Y. Circadian rhythm measurements were performed by N.S.H.E and İ.H.K. All authors contributed the manuscript writing. S.G. edited the manuscript. All the authors read and approved the manuscript.

## Competing interests

The authors declare no competing financial interest.

## Notes

### Competing Interest Statement

The authors have declared no competing interest.

